# Associations Between Vocal Emotion Recognition and Socio-emotional Adjustment in Children

**DOI:** 10.1101/2021.03.12.435099

**Authors:** Leonor Neves, Marta Martins, Ana Isabel Correia, São Luís Castro, César F. Lima

## Abstract

The human voice is a primary channel for emotional communication. It is often presumed that being able to recognise vocal emotions is important for everyday socio-emotional functioning, but direct empirical evidence for this remains scarce. Here, we examined relationships between vocal emotion recognition and socio-emotional adjustment in children. The sample included 6 to 8-year-old children (*N* = 141). The emotion tasks required them to categorise five emotions conveyed by nonverbal vocalisations (e.g., laughter, crying) and speech prosody: anger, disgust, fear, happiness, sadness, plus neutrality. Socio-emotional adjustment was independently evaluated by the children’s teachers using a multi-dimensional questionnaire of self-regulation and social behaviour. Based on frequentist and Bayesian analyses, we found that higher emotion recognition in speech prosody related to better general socio-emotional adjustment. This association remained significant even after accounting for the children’s general cognitive ability, age, sex, and parental education in multiple regressions. Follow-up analyses indicated that the advantages were particularly robust for the socio-emotional dimensions prosocial behaviour and cognitive and behavioural self-regulation. For emotion recognition in nonverbal vocalisations, no associations with socio-emotional adjustment were found. Overall, these results support the close link between children’s emotional prosody recognition skills and their everyday social behaviour.

## Introduction

We perceive emotional information through multiple communication channels, including vocal and facial expressions. These channels offer a window into the emotions of others in social interactions, and the ability to recognise the conveyed emotions may be pivotal for skillful communication. Most research has focused on facial expressions. Nonetheless, the human voice is a major source of emotional information that reflects a primitive and universal form of communication (Ghazanfar & Rendall, 2008; Latinus & Belin, 2011). For instance, hearing a scream of fear can make us realise that someone needs help, or that there is a threat that we might need to attend to. Nonverbal emotional cues in the human voice include inflections in speech, so-called emotional prosody, and purely nonverbal vocalisations, such as laughter and crying.

Emotional prosody corresponds to suprasegmental and segmental modifications in spoken language during emotion episodes. Prosodic cues include pitch, loudness, *tempo,* rhythm, and timbre (Grandjean et al., 2006; Schirmer & Kotz, 2006). Nonverbal vocalisations such as laughter do not contain any linguistic information, and they represent a more primitive form of communication that has been described as the auditory equivalent of facial expressions (Belin et al., 2004). Prosody and nonverbal vocalisations rely on partly distinct articulatory and perceptual mechanisms (Pell et al., 2015; Scott et al., 2010). Based primarily on studies with adult samples, we know that listeners can accurately identify a range of positive and negative emotions from the two types of vocal emotional cues, even when they are heard in isolation and without contextual information (e.g., Castro & Lima, 2010; Cowen et al., 2019; Lima et al., 2013a; Sauter et al., 2010). But it has also been shown that emotion recognition accuracy is higher for nonverbal vocalisations than for emotional prosody (Hawk et al., 2009; Kamiloglu et al., 2020; Sauter et al., 2013).

From a developmental standpoint, soon after birth, infants are able to discriminate emotional expressions in nonverbal vocalisations (e.g., Soderstrom et al., 2017) and prosodic cues (e.g., Flom & Bahrick, 2007). Emotion recognition abilities improve throughout childhood and adolescence, although it is still not established when they peak (Amorim et al., 2019; Grossmann et al., 2010; Morningstar et al., 2018; Sauter et al., 2013). Infants and young children also show a general preference for auditory over visual information (e.g., tones *vs*. lights, Nava & Pavani, 2013; natural sounds *vs*. pictures, Wille & Ebersbach, 2016), which might extend to emotional cues. For instance, Ross et al. (2021) have recently found that children under the age of 8 find it challenging to ignore vocal emotional cues in multimodal stimuli, even if explicitly asked to base their judgment on body cues alone.

Even though it is typically assumed that vocal emotion recognition skills are important for communication at any age, the existing literature has mainly focused on more basic acoustic, perceptual and neurocognitive aspects of these expressions (e.g., Grandjean, 2020; Schirmer & Kotz, 2006). Evidence for associations between vocal emotions and broader aspects of everyday socio-emotional functioning remains relatively scarce, particularly in normative samples. Socio-emotional functioning has been defined as a multidimensional and relatively broad concept (Edwards & Denham, 2018). It includes the ability to accurately understand our own and others’ emotional states, to regulate our own behaviour, and to establish and maintain relationships (Denham et al., 2015; Murray et al., 2015). These processes start to develop early in life and are linked to health outcomes and well-being, both in adults (Nelis et al., 2011) and children (Ogren & Johnson, 2020).

Studies on clinical populations are suggestive of a link between vocal emotional processing and socio-emotional functioning. For instance, neural and behavioural responses to laughter are reduced in boys at risk for psychopathy, who present with disruptive antisocial behaviour and impaired social connectedness (O’Nions et al., 2017). Impairments in the recognition of emotional prosody are further reported in clinical conditions in which deficits in social functioning prevail, namely in individuals at high risk for psychosis and first-episode schizophrenia (Amminger et al., 2012). In the same vein, Parkinson’s disease patients also present impaired emotional prosody recognition (e.g., Lima et al., 2013b), and their atypical prosody during speech production can negatively impact how they are perceived by others (Jaywant & Pell, 2009). There are fewer studies focused on healthy samples, but they point in the same direction. Carton et al. (1999) showed that better emotional prosody recognition was associated with better self-reported relationship well-being in healthy adults, even after controlling for depressive symptoms. Terracciano et al. (2003) also found that better emotional prosody recognition correlated with self-reported openness to experience, a trait linked to social behaviour engagement (e.g., Cabrera et al., 2006; Saef et al., 2018). We have recently shown that the ability to recognise laughter authenticity is associated with higher empathic concern and trait emotional contagion in adults (Neves et al., 2018). However, there are also null results regarding vocal emotion recognition and traits associated with social behaviour, such as agreeableness and extraversion (Furnes et al., 2019).

Children, like adults, make use of vocal emotions in social interactions, and it is important to understand how this relates to their socio-emotional adjustment, given that childhood is a pivotal period for socio-emotional development (Edwards & Denham, 2018; Denham et al., 2015). Studies with preschoolers found that higher emotional prosody recognition correlates with higher peer-rated popularity and lower teacher-rated emotional/behavioural problems (Nowicki & Mitchell, 1988), as well as with lower parent-rated hyperactivity and conduct problems (Chronaki et al., 2015). Studies with school-age children have also documented associations between emotional prosody recognition and socio-emotional variables including self-reported social avoidance and distress (McClure & Nowicki, 2001), teacher-rated social competence (Leppänen & Hietanen, 2001) and emotional and behavioural difficulties (Nowicki et al., 2019), and peer-rated popularity (Leppänen & Hietanen, 2001). However, some of the identified associations are limited to a particular group (e.g., observed for girls, but not for boys; Leppänen & Hietanen, 2001; Nowicki & Mitchell, 1998), and null results have been reported. For instance, preschoolers’ emotional prosody recognition did not correlate with teacher-rated externalising problems (Nowicki & Mitchell, 1998) and parent-rated internalising behaviour (Chronaki et al., 2015). Additionally, inferences have often been based on relatively small samples, typically including less than 80 children, and the focus has been on prosody, leaving nonverbal vocalisations unexplored. To our knowledge, only one study included nonverbal vocalisations, and the emphasis was on how children matched vocal with facial information (Scheerer et al., 2020). Other poorly understood questions are whether associations between vocal emotion recognition and socio-emotional functioning can be seen over and above general differences in cognitive abilities and socio-economic background. These are known to correlate with both emotion recognition (e.g., Erhart et al., 2019; Izard et al., 2000) and social functioning (e.g., Bellanti & Bierman, 2000; Dearing et al., 2006; Gilman et al., 2003), and are often not considered as potential confounds.

In the current study, we asked whether vocal emotion recognition relates to socio-emotional adjustment in 6 to 8-year-old children. We focused on both emotional speech prosody and nonverbal vocalisations and hypothesized that higher emotion recognition accuracy would be associated with better socio-emotional functioning. If children with a greater ability to recognise emotions from vocal cues are better at interpreting social information, this could favor everyday socio-emotional functioning outcomes, such as the willingness to be friendly and helpful with others (prosociality), and the ability to stay calm and focused (self-regulation). Participants completed forced-choice emotion recognition tasks focused on the two types of vocal emotional cues. Their teachers were asked to evaluate children’s socio-emotional functioning using a recently developed multidimensional measure, The Child Self-Regulation and Behaviour Questionnaire (CSBQ; Howard & Melhuish, 2017). This measure has been shown to correlate with outcomes such as peer relationship problems and emotional symptoms (Howard & Melhuish, 2017). We predicted that children scoring higher on vocal emotion recognition would be rated by their teachers as more socio-emotionally competent. We also expected this association to remain significant even when individual differences in age, sex, parental education, and cognitive ability are accounted for. More exploratory questions asked which socio-emotional functioning dimensions are more clearly linked to vocal emotion recognition, and whether associations between emotion recognition and social-emotional functioning are specific to the auditory domain, or are similarly seen across sensory modalities. To that end, children completed an additional emotion recognition task focused on facial expressions. There is some evidence that better facial emotion recognition relates to less behavioural problems (Chronaki et al., 2015; Nowicki & Mitchell, 1998; Nowicki et al., 2019) and better self-regulation skills in children (Rhoades et al., 2009; Salisch et al., 2015), but null results have also been reported, namely regarding social avoidance and distress (McClure & Nowicki, 2001), and peer popularity (Leppänen & Hietanen, 2001).

## Method

### Participants

One hundred forty-eight children were recruited from elementary public schools in a metropolitan area in Northern Portugal (Porto). Seven were excluded due to neurological diseases (*n* = 2), atypically low general cognitive ability (Ravens’ score < 25th percentile; *n* = 4), or lack of data regarding the socio-emotional measure (*n* = 1). The final sample included 141 children (73 boys) between 6 and 8 years of age (*M* = 7.14 years, *SD* = 0.51, range = 6.34 - 8.89). They were 2^nd^ graders from 7 different classes, each of which had one teacher assigned for the entire year. All children were Portuguese native speakers and had normal hearing according to parent reports. Parents’ education varied from 4 to 19 years (*M* = 10.98; *SD* = 3.46). Participants were tested as part of a wider longitudinal project looking at the effects of music training on emotion recognition and socio-emotional behaviour.

An *a priori* power analysis with G*Power 3.1 (Faul et al., 2009) indicated that a sample size of at least 138 would be required to detect correlations of *r* = .30 or larger between variables, considering an alpha level of .05 and a power of .95. For regression models including five predictors (age, sex, parental education, general cognitive ability, and emotion recognition), a sample of at least 134 participants would be required to detect partial associations of *r* = .30 or larger between each predictor variable and socio-emotional adjustment.

This study was approved by the local ethics committee, ISCTE – University Institute of Lisbon (reference 28/2019), and it was conducted in agreement with the Declaration of Helsinki. Written informed consent was obtained for all participants from a parent or legal guardian, and children gave verbal assent to participate.

## Materials

### Emotion Recognition Tasks

The children completed three emotion recognition tasks. Two of them were focused on vocal emotions, speech prosody and nonverbal vocalisations, and the third one on facial expressions. Each task included 60 trials, with 10 different stimuli for each of the six emotional categories (anger, disgust, fear, happiness, sadness, and neutrality). The stimuli were part of previously validated corpora (speech prosody, Castro & Lima, 2010; nonverbal vocalisations, Lima et al., 2013a; facial expressions, Karolinska Directed Emotional Faces database, Goeleven et al., 2008) that have been frequently used (e.g., Agnoli et al., 2012; Correia et al., 2019, 2020; Lima & Castro, 2011; Lima et al., 2016; Lima et al., 2013b; Safar & Moulson, 2020). Speech prosody stimuli were short sentences (*M* = 1473 ms, *SD* = 255) with emotionally neutral semantic content (e.g., “O quadro está na parede”, *The painting is on the wall),* produced by two female speakers to communicate emotions with prosodic cues alone. Nonverbal vocalisations consisted of brief vocal sounds (*M* = 966 ms, *SD* = 259) without linguistic content, such as laughs, screams, or sobs, and were produced by two adult female and two adult male speakers. Facial expressions consisted of colour photographs of male and female actors without beards, moustaches, earrings, eyeglasses, or visible make-up. Each photograph remained visible until participants responded. Based on validation data from adult samples, the average recognition accuracy for the stimuli used in this study was expected to be high (emotional prosody: 78.42%; nonverbal vocalisations: 82.20%; facial expressions: 82.98%).

Participants made a six-alternative forced-choice for each stimulus in each of the three tasks. They were asked to identify the expressed emotion from a list that included *neutrality, anger, disgust, fear, happiness, and sadness*. To improve children’s engagement throughout the task, an emoji illustrating each emotional category was included both on the response pad and on the laptop screen (visible after the presentation of each stimulus). Each task started with six practice trials (one per emotional category), during which feedback was given. After these, the stimuli were presented randomly across two blocks of 30 trials each (no feedback was given). Short pauses were allowed between blocks to ensure that children remained focused and motivated. Each task took approximately 12 minutes. The tasks were implemented using SuperLab Version 5 (Cedrus Corporation, San Pedro, CA), running on an Apple MacBook Pro laptop. Responses were collected using a seven-button response pad (Cedrus RB-740). Auditory stimuli were presented via headphones (Sennheiser HD 201).

The percentage of correct answers was calculated for each emotional category and task. Accuracy rates were corrected for response biases using unbiased hit rates, or *Hu* (Wagner, 1993; for a discussion of biases in forced-choice tasks see, e.g., Isaacowitz et al., 2007). Hu values represent the joint probability that a given emotion will be correctly recognised (given that it is presented), and that a given response category will be correctly used (given that it is used at all), such that they vary between 0 and 1. Hu = 0 when no stimulus from a given emotion is correctly recognised, and Hu = 1 when all the stimuli from a given emotion are correctly recognised (e.g., sad prosody), and the corresponding response category (sadness) is always correctly used (i.e., when there are no false alarms). Primary analyses were conducted using average scores for each task because we had no predictions regarding specific emotions.

### Socio-emotional Adjustment

The Child Self-Regulation and Behaviour Questionnaire (CSBQ) is a 33-item educator-report (or parent-report) questionnaire that assesses children’s socio-emotional behaviour (Howard & Melhuish, 2017). Scale items cover seven subscales: sociability (7 items, e.g., *Chosen as a friend by others),* externalising problems (5 items, e.g., *Aggressive to children),* internalising problems (5 items, e.g., *Most days distressed or anxious),* prosocial behaviour (5 items, e.g., *Plays easily with other children),* behavioural self-regulation (6 items, e.g., *Waits their turn in activities),* cognitive self-regulation (5 items, e.g., *Persists with difficult tasks),* and emotional self-regulation (6 items, e.g., *Is calm and easy going*). Items are rated on a scale from 1 (*not true*) to 5 (*certainly true*). Individual item scores are then summed to produce total scores for each subscale (Howard & Melhuish, 2017). A global socio-emotional functioning score was also computed by averaging the means of the seven subscales, hereafter referred to as *general socio-emotional index*. For this purpose, scores for the externalising and internalising problems subscales were reversed so that higher scores indicated better socio-emotional adjustment across all subscales.

The CSBQ translation to European Portuguese followed the guidelines for adapting tests into multiple languages (e.g., Hambleton, 2005). Two European Portuguese native speakers independently translated the items of the original English CSBQ. They were fluent in English and one of them (C.F.L.) is experienced in the adaptation of questionnaires, and an expert in emotion processing. A single version of the questionnaire was obtained by sorting out the disagreements between the two translators. This version was then shown to two lab colleagues for a final check on language clarity and naturalness, and to discuss the matching between the original and the translated version.

The original CSBQ has sound psychometric properties (Howard & Melhuish, 2017), and in the current dataset internal consistency values were good-to-excellent (Cronbach’s α = 0.85 for general socio-emotional index, ranging from α = 0.80 for externalising/internalising problems to α = 0.91 for cognitive self-regulation).

### General Cognitive Ability

The Raven’s Coloured Progressive Matrices were used as a measure of general nonverbal cognitive ability (Raven, 1947). All participants of the final sample performed within the normative range (≥ 14 out of 36, *M* = 22.63, *SD* = 4.53, range = 14 – 33; norms for Portuguese 2^nd^ graders; Simões, 1995).

#### Procedure

Children were tested individually, in two experimental sessions lasting about 45 minutes in total, in a quiet room at their school. General cognitive ability was assessed in the first session, and emotion recognition in the second one. The order of the emotion recognition tasks was counterbalanced across participants. Before the sessions, a parent completed a background questionnaire that asked for information about parental education and employment, and the child’s history of health issues, such as psychiatric, neurological and hearing impairments.

The CSBQ questionnaire was completed by the children’s teacher. The teachers were blind to the hypothesis of the study, and they had known the children for about one and a half years when they filled the questionnaire.

#### Data Analysis

The data were analysed using standard frequentist *and* Bayesian approaches (e.g., Jarosz & Wiley, 2014). In addition to *p* values, a Bayes Factor (BF_10_) statistic was estimated for each analysis. Bayes factors consider the likelihood of the observed data given the alternative and null hypotheses. The analyses were conducted on JASP Version 0.14.1 (JASP Team, 2020), using the default priors (correlations, stretched beta prior width = 1; *t*-tests, zero-centred Cauchy prior with scale parameter 0.707; linear regressions, JZS prior of *r* = .354; repeated-measures ANOVAs, zero-centered Cauchy prior with a fixed-effects scale factor of *r* = .5, a random-effects scale factor of *r* = 1, and a covariates scale factor of *r* = .354). BF_10_ values were interpreted according to Jeffreys’ guidelines (Jarosz & Wiley, 2014; Jeffreys, 1961). Values below 1 correspond to evidence in favor of the null hypothesis: values between 0.33 and 1 correspond to anecdotal evidence, between 0.10 and 0.33 to substantial evidence, between 0.03 and 0.10 to strong evidence, between 0.01 and 0.03 to very strong evidence, and less than 0.01 to decisive evidence. Values above 1 correspond to evidence for the alternative hypothesis: values between 1 and 3 correspond to anecdotal evidence, between 3 and 10 to substantial evidence, between 10 and 30 to strong evidence, between 30 and 100 to very strong evidence, and greater than 100 to decisive evidence. An advantage of Bayesian statistics is that they allow us to interpret null results and to draw inferences based on them.

For frequentist analyses, Holm-Bonferroni corrections for multiple comparisons were applied whenever appropriate.

The full data set can be found here: https://osf.io/qfp83/?view_only=47031990843a48978ca8058e98118805.

## Results

### Emotion Recognition

Figure 1 shows children’s accuracy in the emotion recognition tasks (see Supplementary Table S1 for statistics for each emotion). Average scores were .41 for speech prosody (*SD* = .18; range = .04 – .85), .72 for vocalisations (*SD* = .11; range = .35 – .94), and .67 for faces (*SD* = .13; range .35 – .94). Performance was above the chance level (.17) for all three modalities, *p*s < .001, BF_10_ > 100, and there was no substantial departure from normality (skewness, range = −1.38 – 0.75; kurtosis, range = −1.36 − 2.64; Curran et al., 1996). Performance differed significantly across tasks, *F*(2, 280) = 296.48, *p* < .001, *η^2^* = .68; BF_10_ > 100. It was lowest for prosody (prosody vs. vocalisations, *p* < .001, BF_10_ > 100; prosody vs. faces, *p* < .001, BF_10_ > 100) and highest for vocalisations (vocalisations vs. faces, *p* < .001, BF_10_ > 100). There was a positive correlation between the two vocal emotion recognition tasks (*r* = .32, *p* < .001, BF_10_ > 100), and between these and the faces task (prosody and faces, *r* = .40, *p* < .001, BF_10_ > 100; vocalisations and faces, *r* = .32, *p* < .001, BF_10_ > 100).

**Figure 1.**
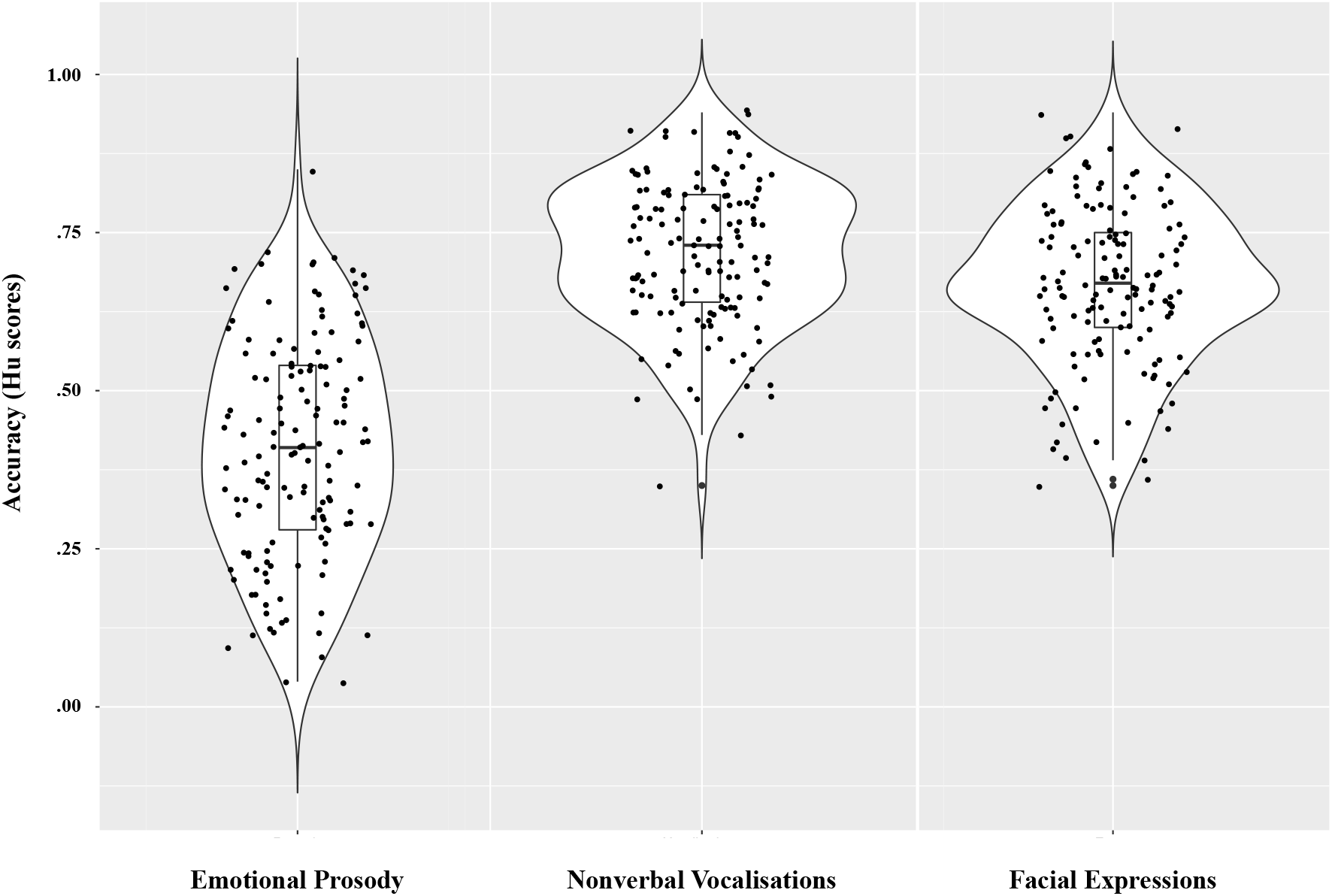
Individual results, box plots and violin plots depicting average emotion recognition scores (Hu) for emotional prosody, nonverbal vocalisations, and facial expressions.

### Socio-emotional Adjustment

Scores for the general socio-emotional index and for each CSBQ subscale are presented in Figure 2. The general socio-emotional score was 3.75 on average, and it varied widely among children, from 2.27 to 4.85 (*SD* = 0.55). There was no substantial departure from normality in the CSBQ data (skewness, range = −0.63 – 0.86; kurtosis, range = −0.84 – 0.05; Supplementary Table S2; Curran et al., 1996). There were correlations among the CSBQ subscales (see Supplementary Table S3 and S4), as expected according to the published data (Howard & Melhuish, 2017).

**Figure 2.**
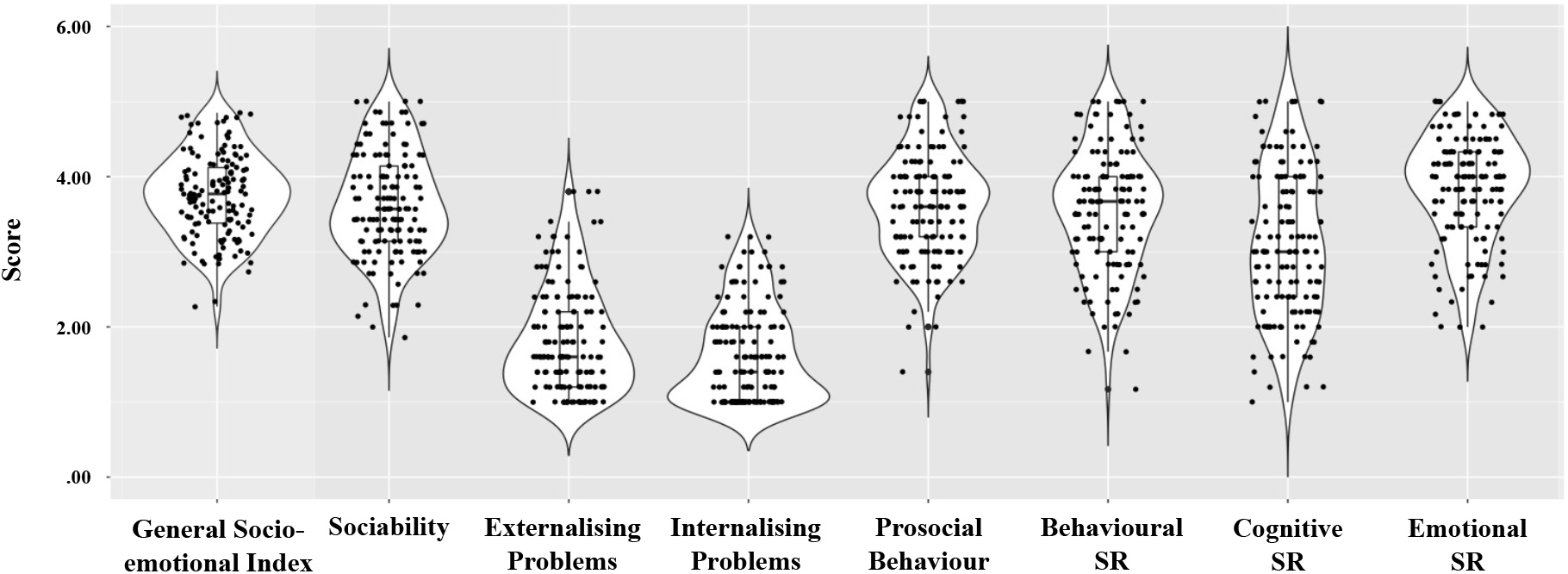
Individual results, box plots and violin plots depicting teacher reports on children’s social-emotional adjustment, as assessed with the CSBQ questionnaire. SR = Self-regulation.

### Cognitive and Socio-demographic Variables

Table 1 shows associations between the main study variables – emotion recognition and general socio-emotional adjustment – and age, sex, parental education, and cognitive ability. Emotion recognition was not associated with demographic or cognitive variables, except for small correlations between emotional prosody recognition and parental education and cognitive ability. Socio-emotional adjustment was higher for girls compared to boys, and it was also higher for younger children and for those with higher parental education.

**Table 1.** Associations between the main study variables (emotion recognition and general socio-emotional adjustment) and age, sex, parental education, and general cognitive ability.

|  | Age | Sex | Parental Education<br>(years) | Cognitive<br>Ability |
| --- | --- | --- | --- | --- |
| Emotion Recognition |  |  |  |  |
| Emotional Prosody | .00<br>*0.11* | .21<br>*0.18* | .25*<br>*8.05* | .27*<br>*22.14* |
| Nonverbal Vocalisations | .14<br>*0.43* | -.63<br>*0.22* | .10<br>*0.21* | .02<br>*0.11* |
| Facial Expressions | .05<br>*0.13* | -1.97<br>*1.06* | .10<br>*0.22* | .10<br>*0.21* |
| General Socio-emotional Index | -.32*<br>*> 100* | -2.97*<br>*9.45* | .42***<br>*> 100* | .22<br>*3.44* |

Note. $N = 141$ for all analyses, except for those involving parental education, where $n = 139$ . $BF_{10}$ values are indicated in italics. For Age, Parental Education and Cognitive Ability, values represent Pearson correlation coefficients; for Sex, they represent $t$ values (two-tailed independent sample $t$ -tests). \* $p < .05$ ; \*\*\* $p < .001$ (Holm Bonferroni-corrected).

### Emotion Recognition and Socio-emotional Adjustment

In line with our prediction, we found decisive evidence that higher emotion recognition in speech prosody related to better general socio-emotional adjustment, *r* = .32, *p* < .001, BF_10_ > 100. A similar association was not found for emotion recognition in nonverbal vocalisations, however, *r* = .10, *p* = .24. It was also not found for faces, *r* = .12, *p* = .33. For both vocalisations and faces, Bayesian analyses provided substantial evidence for the null hypothesis (vocalisations, BF_10_ = 0.21; faces, BF_10_ = 0.27).

To exclude the possibility that the association between emotional prosody recognition and socio-emotional adjustment was due to cognitive or socio-demographic factors, we used multiple regression. We modelled socio-emotional adjustment scores as a function of age, sex, parental education, cognitive ability, and average accuracy on the emotional prosody recognition task. This model explained 30.77% of the variance, *R* = .58, *F*(5,133) = 13.26, *p* < .001, BF_10_ > 100. Independent contributions were evident for age, partial *r* = -.30, *p* < .001, BF_10_ = 49.10, sex, partial *r* = .22, *p* = .01, BF_10_ = 3.06, and parental education, partial *r* = .28, *p* = .001, BF_10_ = 28.68, but not for cognitive ability, *p* = .34, BF_10_ = 0.17. Crucially, emotional prosody recognition made an independent contribution to the model, partial *r* = .27, *p* = .002, and the Bayesian analysis provided strong evidence for this, BF_10_ = 14.25. We calculated Cook’s values and confirmed that this effect was not explained by extreme data points on the regression model (Cook’s distance *M* = 0.01, *SD* = 0.01, range = 0.00 – 0.07). The partial association between emotional prosody recognition and socio-emotional adjustment is illustrated in Figure 3A.

**Figure 3.**
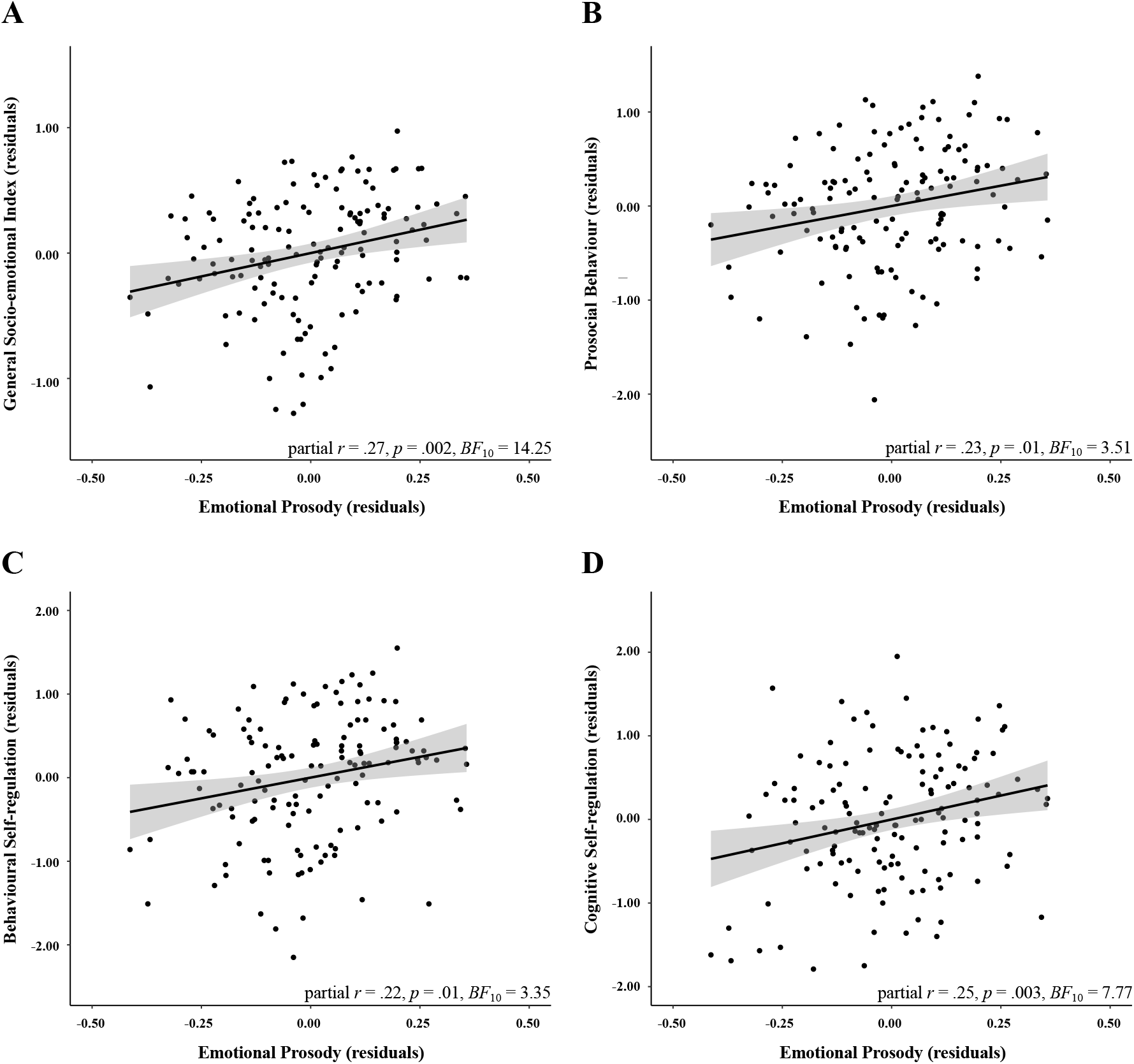
Partial regression plots illustrating the relationship between emotion recognition in speech prosody and general socio-emotional adjustment scores (A), prosocial behaviour (B), behavioural self-regulation (C), and cognitive self-regulation (D), after removing the effects of age, sex, parental education and cognitive ability. Gray shades represent 95% confidence intervals.

Although we had no predictions regarding specific emotions, we wanted to ensure that the association between prosody recognition and socio-emotional adjustment was not driven by a single or small subset of emotions. Follow-up multiple regression analyses, conducted separately for each emotion, showed that positive partial correlations could be seen for all emotions, at significant or trend level (*r*_s_ = .19 – .23), except for disgust (*r* = .13) and sadness (*r* = .12; see Supplementary Table S5 for details).

### Socio-emotional Adjustment Dimensions

We also focused on how emotional prosody recognition related to specific socio-emotional dimensions, considering the CSBQ subscales: sociability, externalising problems, internalising problems, prosocial behaviour, behavioural self-regulation, cognitive selfregulation, and emotional self-regulation. Associations were particularly clear for prosocial behaviour, cognitive self-regulation, and behavioural self-regulation, all supported by substantial evidence (3.34 < BF_10_ < 7.78). This was indicated by multiple regressions modelling scores on each CSBQ subscale as a function of age, sex, parental education, cognitive ability, and average accuracy on emotional prosody recognition. For prosocial behaviour, the model explained 18.60% of the variance, *R* = .46, *F*(5,133) = 7.31, *p* < .001, BF_10_ > 100. Independent contributions were evident for emotional prosody, partial *r* = .23, *p* = .01, BF_10_ = 3.51, sex, partial *r* = .21, *p* = .02, BF_10_ = 2.16, and parental education, partial *r* = .24, *p* = .01, BF_10_ = 4.93, but not for age, *p* = .09, BF_10_ = 0.48, or cognitive ability, *p* = .84, BF_10_ = 0.11. For behavioural self-regulation, the model explained 19.20% of the variance, *R* = .47, *F*(5,133) = 7.56, *p* < .001, BF_10_ > 100. Independent contributions were evident for emotional prosody, partial *r* = .22, *p* = .01, BF_10_ = 3.35, sex, partial *r* = .27, *p* = .001, BF_10_ = 16.34, and parental education, partial *r* = .24, *p* = .01, BF_10_ = 6.11, but not for age, *p* = .26, BF_10_ = 0.21, or cognitive ability, *p* = .83, BF_10_ = 0.11. For cognitive self-regulation, the model explained 42.59% of the variance, *R* = .67, *F*(5,133) = 21.47, *p* < .001, BF_10_ > 100. Independent contributions were evident for emotional prosody, partial *r* = .25, *p* = .003, BF_10_ = 7.77, age, partial *r* = -.23, *p* = .01, BF_10_ = 4.76, parental education, partial *r* = .41, *p* < .001, BF_10_ > 100, and cognitive ability, partial *r* = .31, *p* < .001, BF_10_ > 100, but not for sex, *p* = .54, BF_10_ = 0.13. We calculated Cook’s values and confirmed that these effects were not explained by extreme data points on the regression model: Cook’s distance *M* = 0.01, *SD* = 0.01 (Cook’s distance range = 0.00 – 0.06 for prosocial behaviour; 0.00 – 0.05 for behavioural self-regulation; and 0.00 – 0.06 for cognitive self-regulation). Partial associations between emotional prosody recognition and these dimensions of socio-emotional adjustment are illustrated in Figure 3 B to D.

There were also significant associations between emotional prosody recognition and the dimensions of sociability and emotional self-regulation, but the level of evidence was weaker (1.61 < BF_10_ < 2.74). For sociability, the model explained 18.96% of the variance, *R* = .47, *F*(5,133) = 7.46, *p* < .001, BF_10_ > 100. Independent contributions were evident for emotional prosody, partial *r* = .20, *p* = .02, BF_10_ = 1.62, age, partial *r* = -.33, *p* < .001, BF_10_ > 100, and parental education, partial *r* = .18, *p* = .04, BF_10_ = 0.91, but not for sex, *p* = .79, BF_10_ = 0.11, or cognitive ability, *p* = .49, BF_10_ = 0.14. For emotional self-regulation, the model explained 9.26% of the variance, *R* = .35, *F*(5,133) = 3.82, *p* = .003, BF_10_ = 5.05. Independent contributions were evident for emotional prosody, partial *r* = .22, *p* = .01, BF_10_ = 2.73, and sex, partial *r* = .23, *p* = .01, BF_10_ = 4.34, but not for age, *p* = .15, BF_10_ = 0.30, parental education, *p* = .53, BF_10_ = 0.13, or cognitive ability, *p* = .15, BF_10_ = 0.29.

For the remaining two socio-emotional dimensions, externalising and internalising problems, emotional prosody recognition did not uniquely contribute to the models (*p*s > .33, BF_10_ < 0.18).

## Discussion

In the current study, we asked whether individual differences in vocal emotion recognition relate to socio-emotional adjustment in children. We measured emotion recognition in two types of vocal emotions, speech prosody and nonverbal vocalisations. Socio-emotional adjustment was assessed through a multidimensional measure completed by the children’s teachers. We found strong evidence for a positive association between speech prosody recognition and socio-emotional adjustment, based on both frequentist and Bayesian statistics. This association remained significant even after accounting for age, sex, parental education, and cognitive ability. Follow-up analyses further showed that prosody recognition was more robustly linked to the socio-emotional dimensions prosocial behaviour, cognitive self-regulation, and behavioural self-regulation. For emotion recognition in nonverbal vocalisations, there were no associations with socio-emotional adjustment. A similar null result was found for the additional emotion recognition task focused on facial expressions.

Some prior studies have reported an association between children’s emotional prosody recognition abilities and aspects of socio-emotional adjustment including behavioural problems (e.g., social avoidance and distress; McClure & Nowicki, 2001), peer popularity (e.g., Nowicki & Mitchell, 1998), and global social competence (e.g., Leppänen & Hietanen, 2001). However, results have been mixed (Chronaki et al., 2015; Nowicki & Mitchell, 1998) and often based on relatively small samples. It also remained unclear whether the associations are specific, or a result of factors such as parental education. The present study corroborates the association between emotional prosody recognition and socio-emotional adjustment in a sample of 6 to 8-year-olds, and indicates that this association is not reducible to cognitive or socio-demographic variables, namely age, sex, cognitive ability, and parental education. Emotional prosody cues help us build up a mental representation of other’s emotional states (Grandjean, 2020), and prosody can convey a wide range of complex and nuanced states, such as verbal irony, sarcasm and confidence (Cheang & Pell, 2008; Morningstar et al., 2018; Pell & Kotz, 2021). Interpreting prosodic cues might be challenging, as indicated by evidence (that we replicated) that emotion recognition accuracy is lower for emotional prosody compared to nonverbal vocalisations and facial expressions (e.g., Hawk et al., 2009; Kamiloglu et al., 2020; Sauter et al., 2013). This increased difficulty might be because prosodic cues are embedded in speech, which constrains acoustic variability (Scott et al., 2010). These stimuli are also more complex in that they include both lexico-semantic and prosodic cues, while lexico-semantic information is not present in nonverbal vocalisations and facial expressions. Children with an earlier and more efficient development of this complex ability might therefore be particularly well-equipped to navigate their social worlds.

In analyses focused on specific dimensions of socio-emotional adjustment, we found that children’s ability to recognise emotional prosody was particularly related to prosocial behaviour and cognitive and behavioural self-regulation. Prosociality is associated with positive social behaviours such as cooperation, altruism, and empathy (Jensen, 2016; Lockwood et al., 2014). The ability to recognise fearful facial expressions was found to be linked to adults’ prosocial behaviour (Adolphs & Tusche, 2017; Marsh et al., 2007; Marsh et al., 2014). This could be because distress cues are a powerful tool to elicit care, and being able to ‘read’ them could promote prosocial behaviours, such as helping a child who is crying (Marsh, 2019). Regarding vocal emotions, decreased cooperative behaviour was observed in adults towards partners displaying emotional prosody of anger, fear and disgust (Caballero & Díaz, 2019). However, this was found in a study focused on decisions to cooperate in a social decision-making paradigm, and participants’ ability to recognise emotional prosody was not examined. To our knowledge, the present study is the first to show that emotional prosody recognition is positively linked to prosocial behaviour in school-aged children. It is possible that the ability to accurately interpret the emotional meaning of complex stimuli (such as speech) allows children to more readily deduce when to cooperate, share, or help others, all prosocial behaviours covered by our measure.

Self-regulation includes behavioural and cognitive components, and we found associations with prosody for both. The behavioural component refers to the ability to remain on task, to inhibit behaviours that might not contribute to goal achievement, and to follow socially appropriate rules (Murray et al., 2015). The cognitive component is focused on more top-down processes related to problem-solving, focused attention and self-monitoring, which might support autonomy and task persistence. Prior evidence shows that preschoolers’ recognition of facial expressions correlates with attention processes and behavioural selfregulation (Rhoades et al., 2009; Salisch et al., 2015), but evidence regarding vocal emotion recognition is scant. In view of previous reports that attention might contribute to performance in emotional prosody tasks in adults (e.g., Borod et al., 2000; Lima et al., 2013b), it could have been that children who were more able to focus and remain on task were in a better position for improved performance. However, although we found a correlation between cognitive ability and prosody recognition, thus replicating previous evidence, the association with self-regulation remained significant after cognitive ability was accounted for, making this explanation less likely. Alternatively, because the ability to decode emotional prosody supports a more efficient understanding of communicative messages (e.g., from parents or teachers), this might allow children to more easily understand the tasks they are expected to perform, the rules to follow, and the goals to achieve. Future studies assessing self-regulatory processes in more detail will be important to delineate the sub-processes driving the general associations uncovered here.

Contrasting with the findings for emotional prosody, for nonverbal vocalisations we observed no associations with socio-emotional adjustment. To our knowledge, ours is the first study that systematically considers the two sources of nonverbal vocal emotional cues - prosody and nonverbal vocalisations - in the context of associations with socio-emotional functioning. This matters because, despite being both vocal emotional expressions, they differ in their production and perceptual mechanisms (Pell et al., 2015; Scott et al., 2010), and indeed also seem to differ in their correlates. This null result seems unexpected, considering that nonverbal vocalisations reflect a primitive and universal form of communication (e.g., Sauter et al., 2010), thought to play an important role in social interactions. It could have been that our measures of emotion recognition and socio-emotional adjustment were not sensitive enough to capture the effect. But it could also be that variability in the processing of vocalisations does not play a major role for socio-emotional functioning in typically developing school-age children. Previous results indicate that children as young as 5 years are already highly proficient at recognizing a range of positive and negative emotions in nonverbal vocalisations, with average accuracy approaching 80%, and there is no improvement from 5 to 10 years for most emotions (Sauter et al., 2013). Such proficiency is replicated here, and we also found that the range of individual differences is small when compared to prosody (see Figure 1). This could mean that, for the majority of healthy schoolage children, the ability to recognise nonverbal emotional vocalisations is already high enough for them to optimally use these cues in social interactions, such that small individual variation will not necessarily translate into measurable differences in everyday behaviour. This result will need to be followed-up in future studies, however, to examine whether it replicates across different measures and age groups (e.g., including a broader range of emotions and a more comprehensive assessment of socio-emotional adjustment).

That performance on the additional facial emotion recognition task also did not correlate with socio-emotional adjustment corroborates the findings of some previous studies. McClure and Nowicki (2001) found that 8 to 10-year-old children’s ability to recognise facial expressions was not associated with dimensions of socio-emotional adjustment, namely social avoidance and distress. Leppänen and Hietanen (2001) also reported null results regarding peer popularity in a sample of 7 to 10-year-olds. Moreover, Chronaki et al. (2015) found that preschoolers’ ability to recognise facial expressions was not associated with parent-rated internalising problems. On the other hand, there is evidence that facial emotion recognition can relate to fewer behavioural problems in school-age children (e.g., Nowicki et al., 2019) and to better self-regulation in preschoolers (e.g., Salisch et al., 2015). These discrepancies across studies might stem from differences in samples’ characteristics and measures, and they will be clarified as more research is conducted on this topic. In the current study, based on a relatively large sample informed by power analyses, Bayesian statistics provided in fact evidence for the null hypothesis. In line with our reasoning for nonverbal vocalisations, a tentative explanation is that children’s proficiency at decoding facial emotions at this age is already high, such that the impact of individual variation in everyday life behaviour might be less apparent.

A limitation of the current study is the correlational approach. We provide evidence for an association between emotional prosody recognition and socio-emotional adjustment, but we cannot exclude the possibility that emotional prosody recognition skills are the result, not the cause, of better socio-emotional adjustment. Having more and better social interactions could provide opportunities for children to learn about emotional expressions, and to hone their emotion recognition skills. Future systematic longitudinal research will be needed to establish causality, for example by testing whether an emotion recognition training program leads to improved social interactions. Another limitation is that we used vocal and facial stimuli produced by adults, and it would be interesting to know if similar results would be obtained with stimuli produced by children. Children can accurately recognise vocal expressions produced by participants of any age, but there is also evidence that they might perform better for stimuli produced by children their age (Amorim et al., 2019; Rhodes & Anastasi, 2012; but see McClure & Nowicki, 2001). One last point is that we only used a teacher-report socio-emotional measure. Future work combining different socio-emotional measures, such as parent-report and also performance-based tasks, would allow us to more stringently test these relationships.

In conclusion, the current study shows that emotional speech prosody recognition is associated with general socio-emotional adjustment in children. We also show that this association is not explained by cognitive and socio-demographic variables, and results were particularly robust for the socio-emotional dimensions prosocial behaviour and selfregulation (cognitive and behavioural components). These findings did not generalize to vocal emotional stimuli without linguistic information - nonverbal vocalisations - and were also not seen for facial expressions. Altogether, these results support the notion that emotional speech recognition skills play an important role in children’s everyday social interactions. They also contribute to debates on the functional role of vocal emotional expressions, and might inform interventions aimed at fostering socio-emotional skills in childhood.

## Data Availability

The dataset supporting this article is freely available for public use at the OSF platform: https://osf.io/qfp83/?view_only=47031990843a48978ca8058e98118805

## Competing Interesting Statement

César F. Lima was a member of the Royal Society Open Science editorial board at the time of submission; however, they were not involved in the editorial assessment of the manuscript in any way.

## Research Ethics

Ethical approval for the study protocol was obtained from the local Ethics Committee, ISCTE-IUL (reference 28/2019). Written informed consent was collected from all participants from a parent or legal guardian, and children gave verbal assent to participate.

## Authors’ Contributions

LN and MM participated in the design of the study, prepared the tasks, collected the data, conducted data analysis and interpretation, and drafted the manuscript. AIC collected the data, participated in data analysis, and helped draft the manuscript. SLC helped conceive and design the study, coordinated the study, and helped draft the manuscript. CFL conceived and designed the study, coordinated the study, participated in data analysis and interpretation, and drafted the manuscript. All authors gave final approval for publication.

## Acknowledgments

We thank the school administration, teachers, parents, and all the children who took part in the study.

## Supplementary information

**Supplementary Table S1.** Summary statistics for emotion recognition in emotional prosody, nonverbal vocalisations and facial expressions.

| Emotion recognition<br>(average Hu scores) | Mean ± SD (min – max) | Skewness | Kurtosis |
| --- | --- | --- | --- |
| Emotional Prosody |  |  |  |
| Neutral | .28 ± .19 (.00 – .83) | 0.44 | -0.13 |
| Happy | .52 ± .21 (.01 – 1.00) | -0.18 | -0.35 |
| Sad | .29 ± .23 (.00 – .81) | 0.13 | -1.36 |
| Fear | .48 ± .29 (.00 – 1.00) | -0.16 | -1.17 |
| Angry | .56 ± .28 (.00 – 1.00) | -0.42 | -0.81 |
| Disgust | .31 ± .27 (.00 – 1.00) | 0.75 | -0.35 |
| Total | .41 ± .18 (.04 – .85) | 0.00 | -0.73 |
| Nonverbal Vocalisations |  |  |  |
| Neutral | .74 ± .23 (.00 – 1.00) | -1.01 | 0.54 |
| Happy | .71 ± .17 (.17 – 1.00) | -0.48 | 0.39 |
| Sad | .75 ± .14 (.30 – 1.00) | -0.41 | 0.26 |
| Fear | .64 ± .22 (.00 – 1.00) | -0.85 | 0.36 |
| Angry | .75 ± .16 (.25 – 1.00) | -0.69 | 0.08 |
| Disgust | .72 ± .17 (.23 – 1.00) | -0.40 | -0.07 |
| Total | .72 ± .11 (.35 – .94) | -0.39 | -0.10 |
| Facial Expressions |  |  |  |
| Neutral | .72 ± .14 (.23 – 1.00) | -0.39 | 0.22 |
| Happy | .89 ± .12 (.36 – 1.00) | -1.38 | 2.64 |
| Sad | .63 ± .21 (.10 – 1.00) | -0.46 | -0.38 |
| Fear | .65 ± .22 (.00 – 1.00) | -1.07 | 1.13 |
| Angry | .56 ± .20 (.08 – 1.00) | -0.10 | -0.44 |
| Disgust | .55 ± .23 (.00 – 1.00) | -0.37 | -0.46 |
| Total | .67 ± .13 (.35 – .94) | -0.30 | -0.27 |

**Supplementary Table S2.**
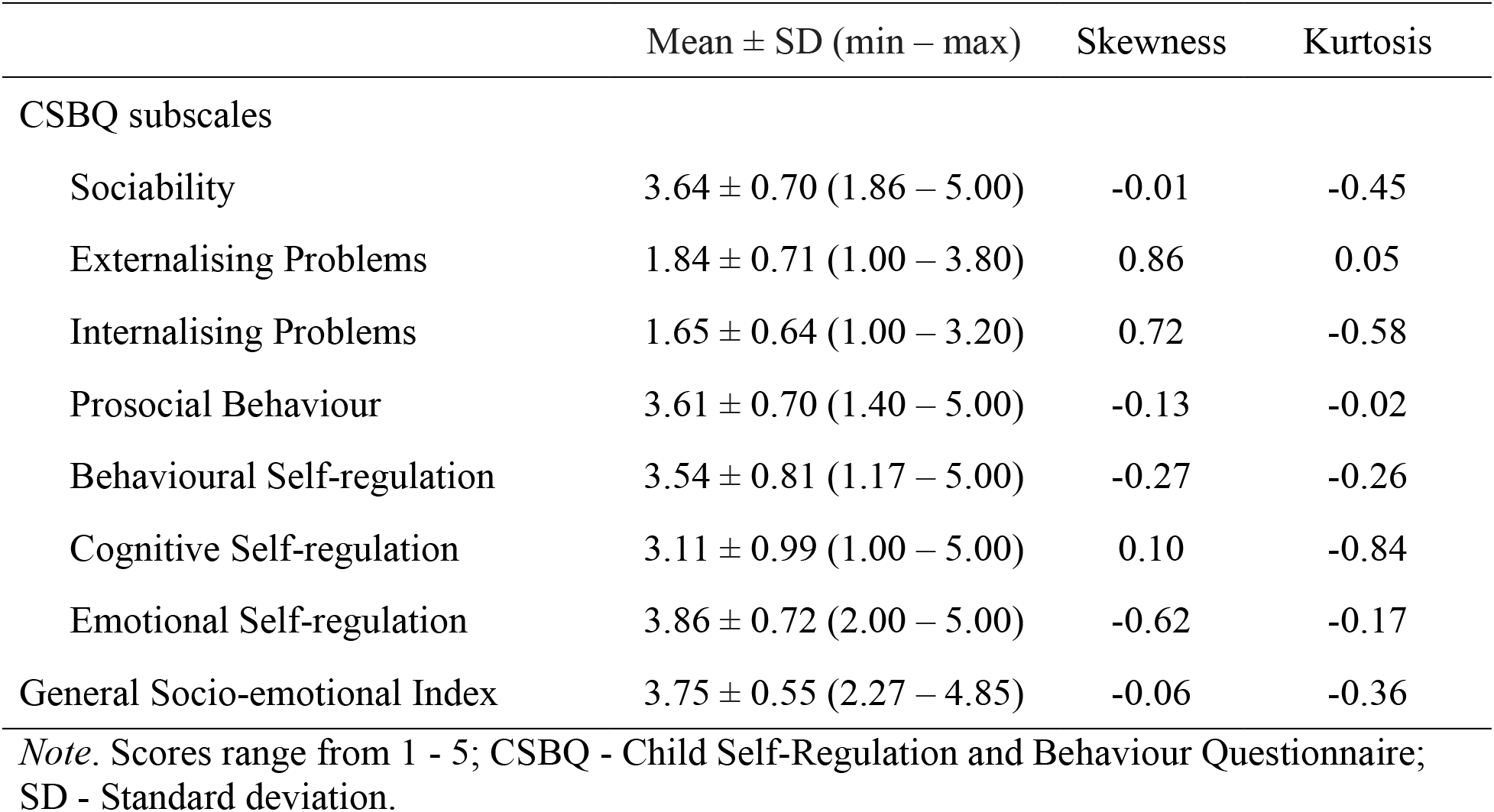
Summary statistics and reliability for the CSBQ subscales (sociability, externalising problems, internalising problems, prosocial behaviour, behavioural self-regulation, cognitive self-regulation,and emotional self-regulation) and general socio-emotional index.

**Supplementary Table S3.**
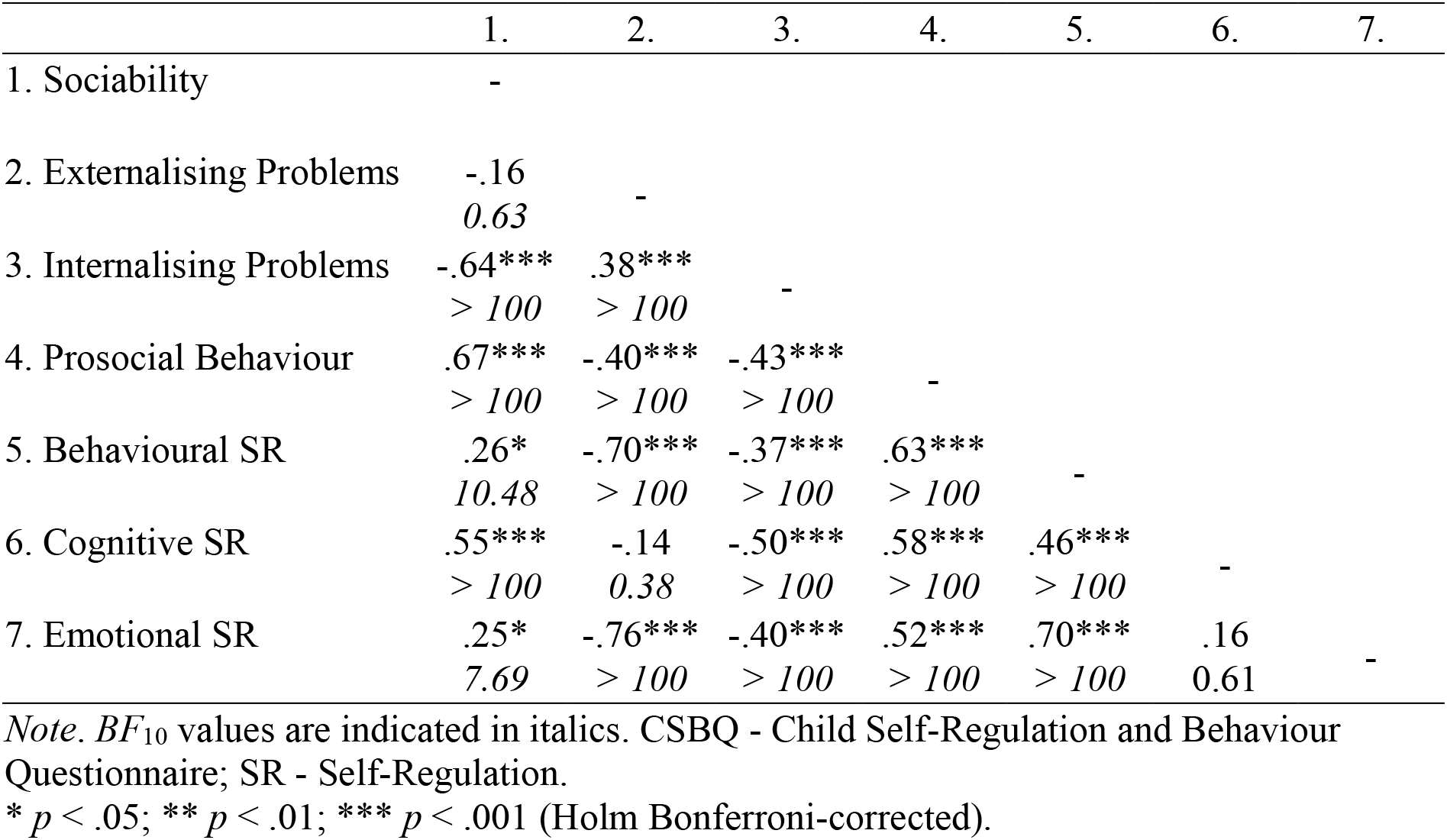
Correlations between the CSBQ subscales.

|  | 1. | 2. | 3. | 4. | 5. | 6. | 7. |
| --- | --- | --- | --- | --- | --- | --- | --- |
| 1. Sociability | - |  |  |  |  |  |  |
| 2. Externalising Problems | -.16<br>*0.63* | - |  |  |  |  |  |
| 3. Internalising Problems | -.64***<br>*> 100* | .38***<br>*> 100* | - |  |  |  |  |
| 4. Prosocial Behaviour | .67***<br>*> 100* | -.40***<br>*> 100* | -.43***<br>*> 100* | - |  |  |  |
| 5. Behavioural SR | .26*<br>*10.48* | -.70***<br>*> 100* | -.37***<br>*> 100* | .63***<br>*> 100* | - |  |  |
| 6. Cognitive SR | .55***<br>*> 100* | -.14<br>*0.38* | -.50***<br>*> 100* | .58***<br>*> 100* | .46***<br>*> 100* | - |  |
| 7. Emotional SR | .25*<br>*7.69* | -.76***<br>*> 100* | -.40***<br>*> 100* | .52***<br>*> 100* | .70***<br>*> 100* | .16<br>*0.61* | - |

*Note.* $BF_{10}$ values are indicated in italics. CSBQ - Child Self-Regulation and Behaviour Questionnaire; SR - Self-Regulation.
\* $p < .05$ ; \*\* $p < .01$ ; \*\*\* $p < .001$ (Holm Bonferroni-corrected).

**Supplementary Table S4.** Pairwise correlations between the general socio-emotional index and each of the CSBQ subscales.

| CSBQ subscales | *r* | BF<sub>10</sub> |
| --- | --- | --- |
| Sociability | .68*** | > 100 |
| Externalising Problems | -.67*** | > 100 |
| Internalising Problems | -.71*** | > 100 |
| Prosocial Behaviour | .83*** | > 100 |
| Behavioural Self-regulation | .81*** | > 100 |
| Cognitive Self-regulation | .70*** | > 100 |
| Emotional Self-regulation | .72*** | > 100 |

Note. CSBQ - Child Self-Regulation and Behaviour Questionnaire.
\*\*\* $p < .001$ (Holm Bonferroni-corrected).

**Supplementary Table S5.** Partial correlations between the general socio-emotional adjustment scores and each category of emotional prosody, after removing the effects of age, sex, parental education and cognitive ability.

| Emotional Prosody Categories | partial $r$ | $p$ | BF<sub>10</sub> | R | Adjusted $R^2$ |
| --- | --- | --- | --- | --- | --- |
| Neutral | .19 | .03 | 1.24 | .56 | .28 |
| Happy | .23 | .01 | 3.81 | .57 | .29 |
| Sad | .12 | .16 | 0.30 | .54 | .27 |
| Fear | .21 | .01 | 2.26 | .56 | .29 |
| Angry | .22 | .01 | 3.19 | .56 | .29 |
| Disgust | .13 | .12 | 0.36 | .54 | .27 |

